## Supplemental Figures and Tables for "Distinct mechanisms of inhibition of Kv2 potassium channels by tetraethylammonium and RY785"

### Supplementary Information

Shan Zhang[1], Robyn Stix[1,2], Esam A. Orabi[1], Nathan Bernhardt[1], José D. Faraldo-Gómez[1*]

[1]Theoretical Molecular Biophysics Laboratory,
National Heart, Lung and Blood Institute,
National Institutes of Health, Bethesda, MD

[2]Molecular and Cell Biology Graduate Program,
Johns Hopkins University, Baltimore, MD

February 24th, 2026

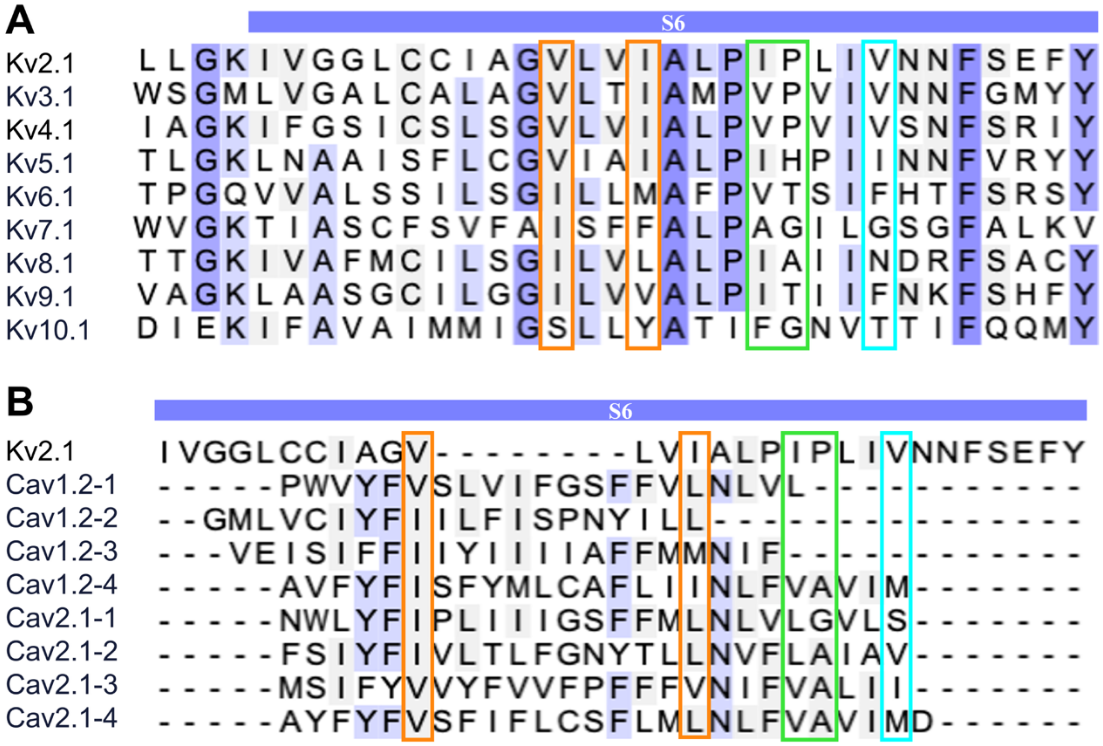


**Supplementary Figure 1.** Multiple-sequence alignment of the S6 segment in selected $K^+$ and $Ca^{2+}$ channels. (**A**) Kv2.1 is compared with other Kv channels. (**B**) Kv2.1 is compared with Cav channels. The degree of sequence similarity is indicated in colors, from dark purple (high) to light purple (low). Orange, green, and cyan rectangles highlight Kv2.1 residues observed to interact with RY785 in our simulations, consistent with the color scheme used in Figure 5. Note that in Cav channels the S6 sequences differ among the four subunits, and thus, each subunit is aligned with Kv2.1 for comparison.

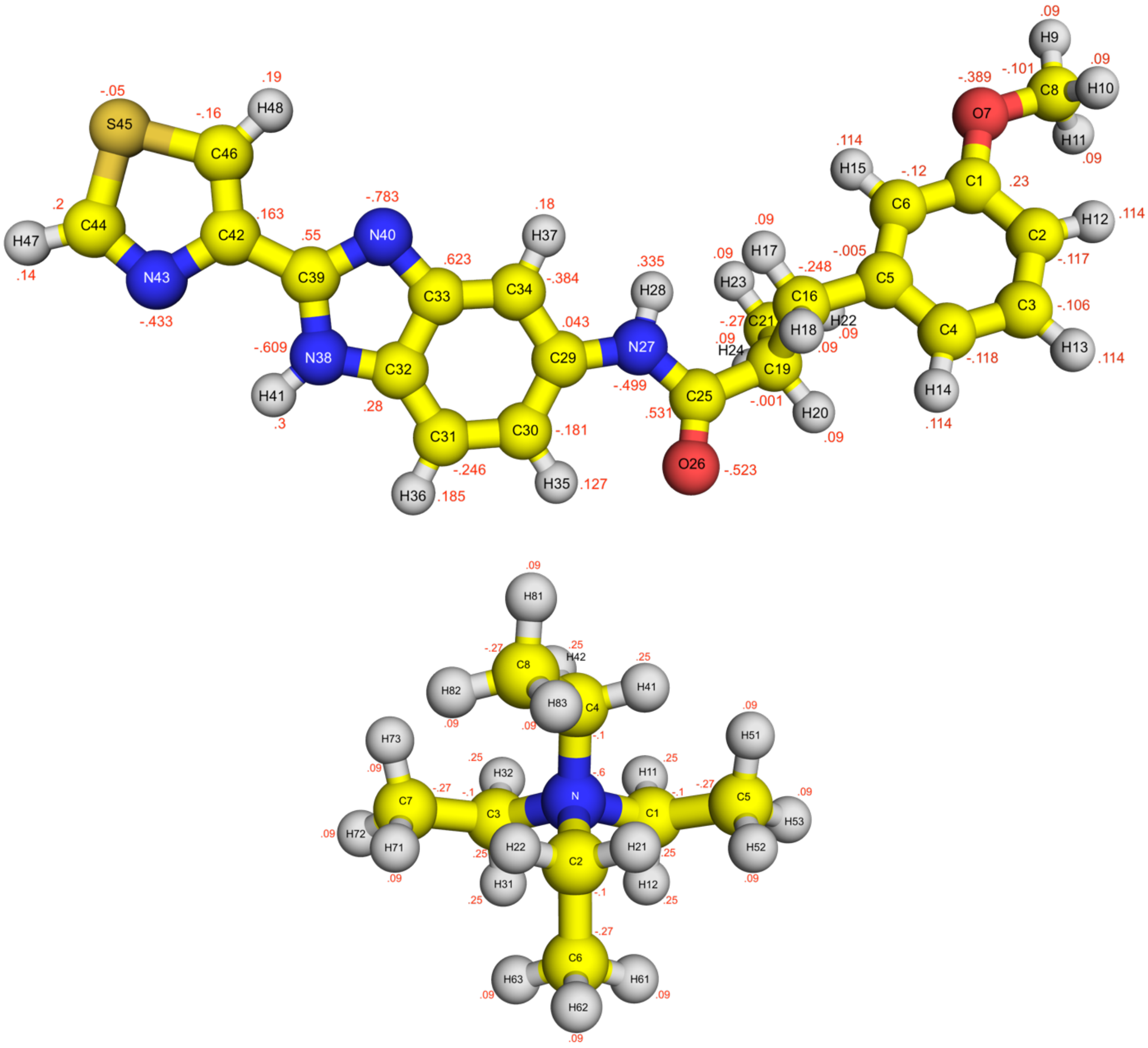


**Supplementary Figure 2.** Chemical structures of (top) RY785 and (bottom) tetraethylammonium. Atom names and partial electronic charges are indicated.

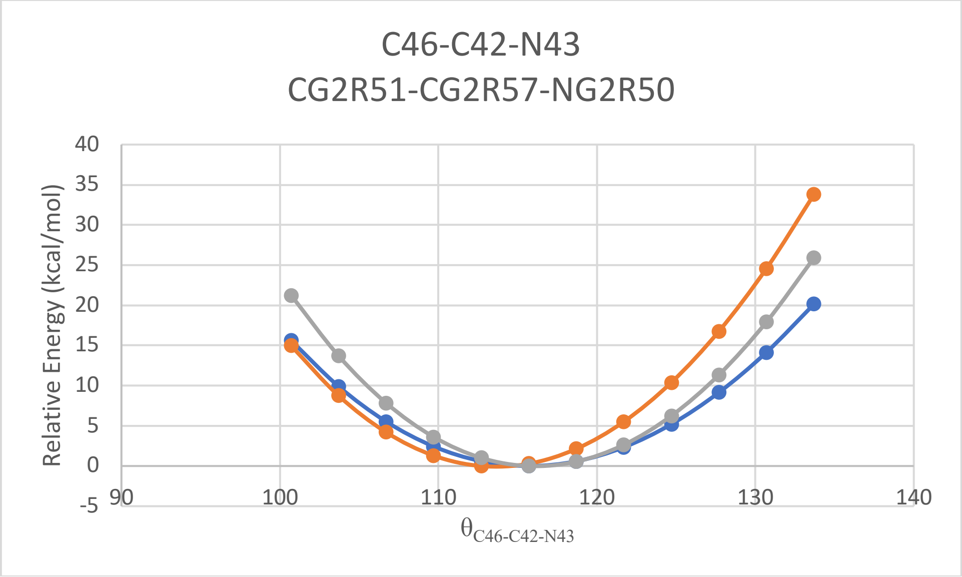


**Supplementary Figure 3.** Potential-energy curve for the C46-C42-N43 bond angle in RY785 between 100° and 135°, calculated with MP2/6-31G(d) (blue) and with default CGenFF (orange) or with our optimized forcefield (gray). The Y axis shows the energy (in kcal/mol) relative to the lowest-energy conformer.

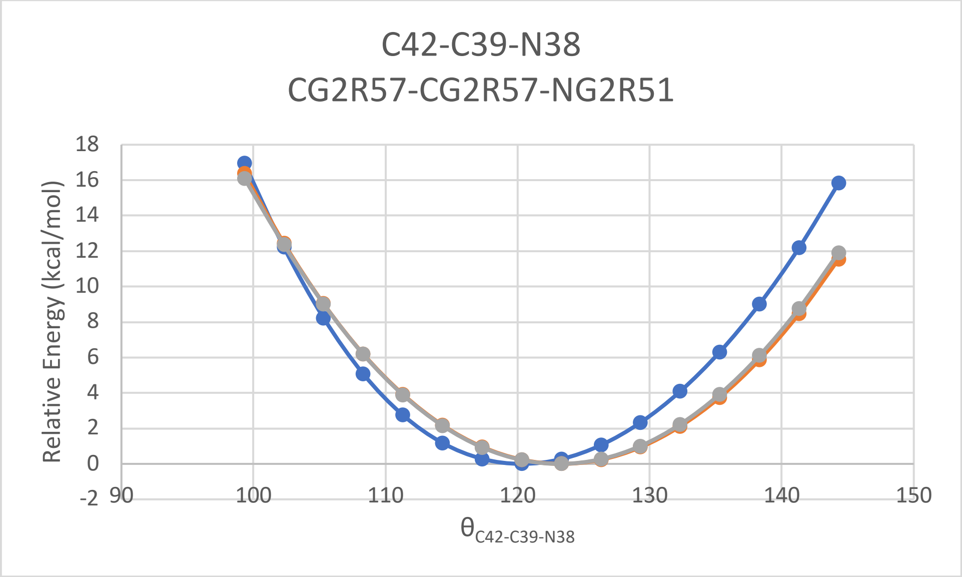


**Supplementary Figure 4.** Potential-energy curve for the C42-C39-N38 bond angle in RY785 between 100° and 145°, calculated with MP2/6-31G(d) (blue) and with default CGenFF (orange) or with our optimized forcefield (gray). The Y axis shows the energy (in kcal/mol) relative to the lowest-energy conformer.

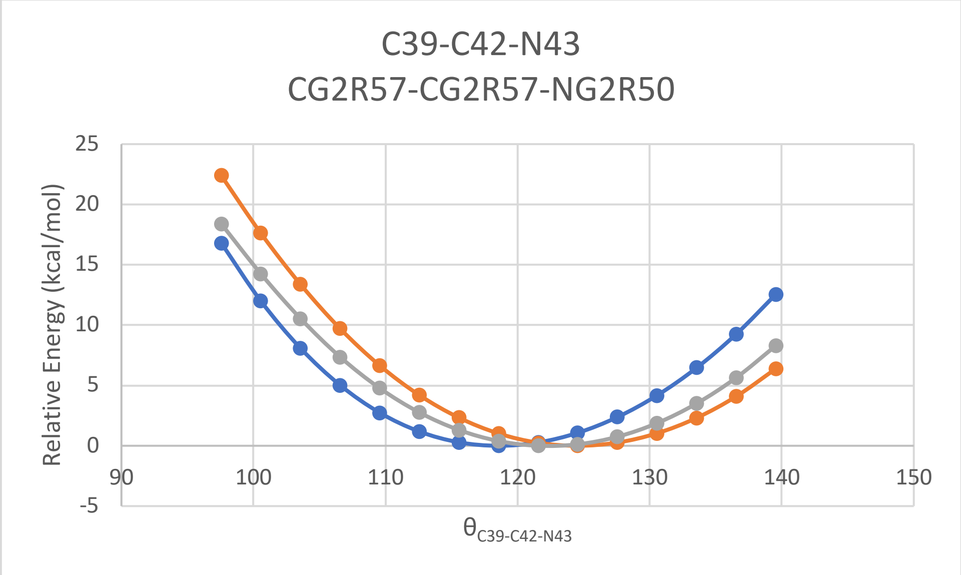


**Supplementary Figure 5.** Potential-energy curve for the C39-C42-N43 bond angle in RY785 between 95° and 140°, calculated with MP2/6-31G(d) (blue) and with default CGenFF (orange) or with our optimized forcefield (gray). The Y axis shows the energy (in kcal/mol) relative to the lowest-energy conformer.

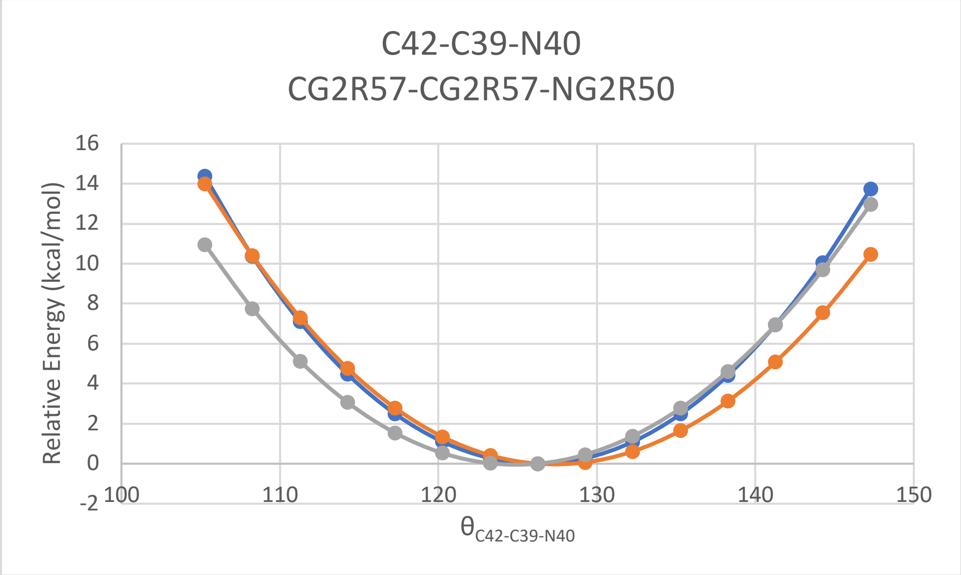


**Supplementary Figure 6.** Potential-energy curve for C42-C39-N40 bond angle in RY785 between 105° and 150°, calculated with MP2/6-31G(d) (blue) and with default CGenFF (orange) or with our optimized forcefield (gray). The Y axis shows the energy (in kcal/mol) relative to the lowest-energy conformer.

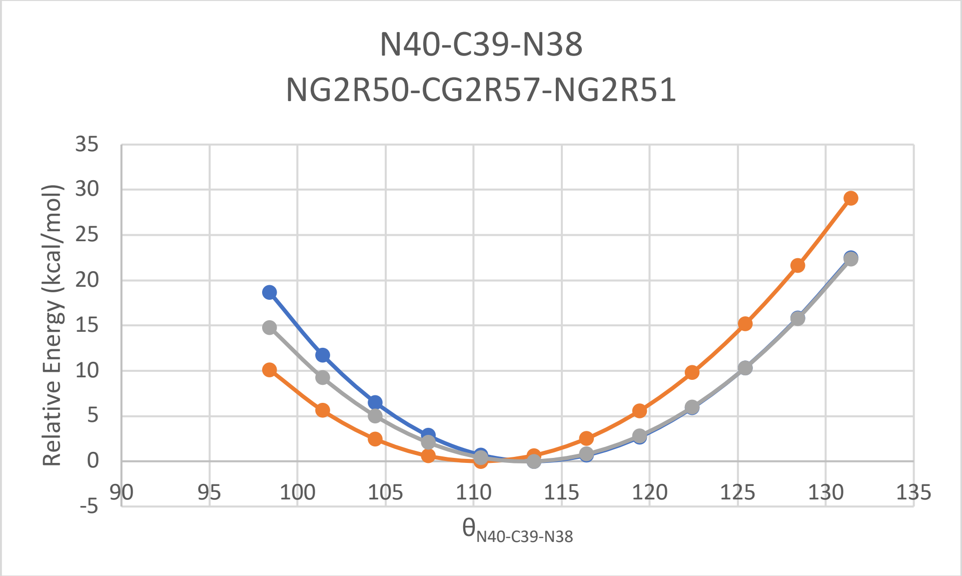


**Supplementary Figure 7.** Potential-energy curve for the N40-C39-N38 bond angle in RY785 between 95° and 135°, calculated with MP2/6-31G(d) (blue) and with default CGenFF (orange) or with our optimized forcefield (gray). The Y axis shows the energy (in kcal/mol) relative to the lowest-energy conformer.

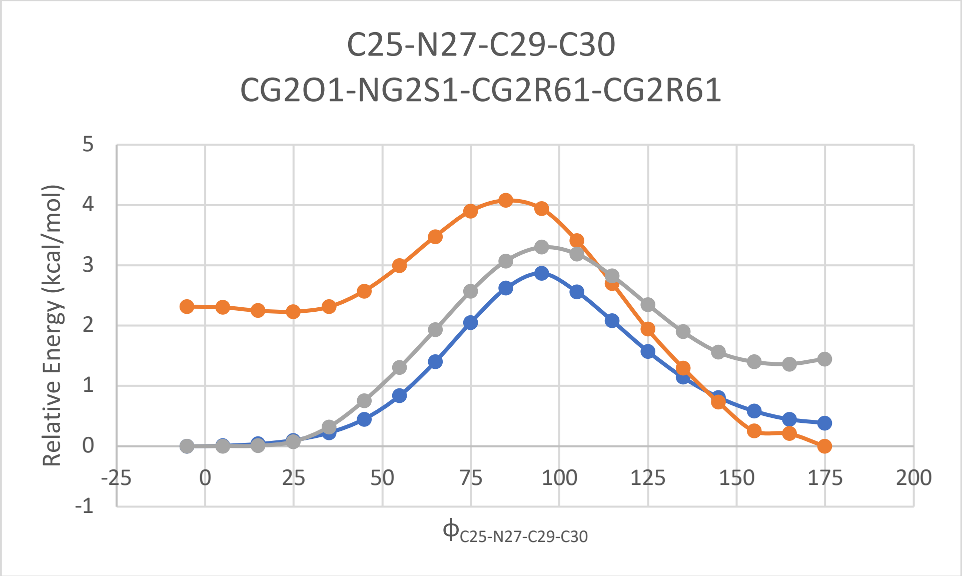


**Supplementary Figure 8.** Potential-energy curve for the C25-N27-C29-C30 dihedral angle in RY785 between 0° and 180°, calculated with MP2/6-31G(d) (blue) and with default CGenFF (orange) or with our optimized forcefield (gray). The Y axis shows the energy (in kcal/mol) relative to the lowest-energy conformer.

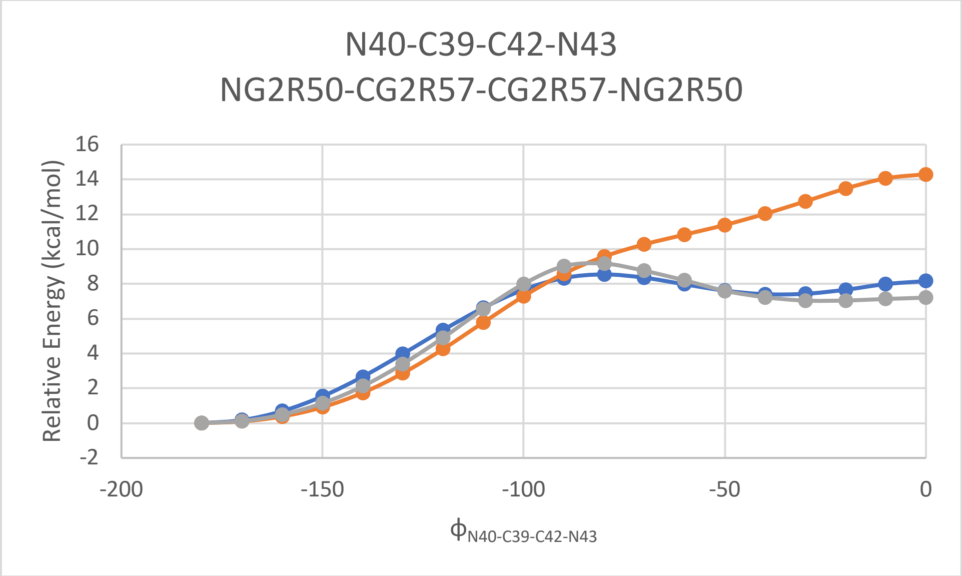


**Supplementary Figure 9.** Potential-energy curve for the N40-C39-C42-N43 dihedral angle in RY785 between -200° and 0°, calculated with MP2/6-31G(d) (blue) and with default CGenFF (orange) or with our optimized forcefield (gray). The Y axis shows the energy (in kcal/mol) relative to the lowest-energy conformer.

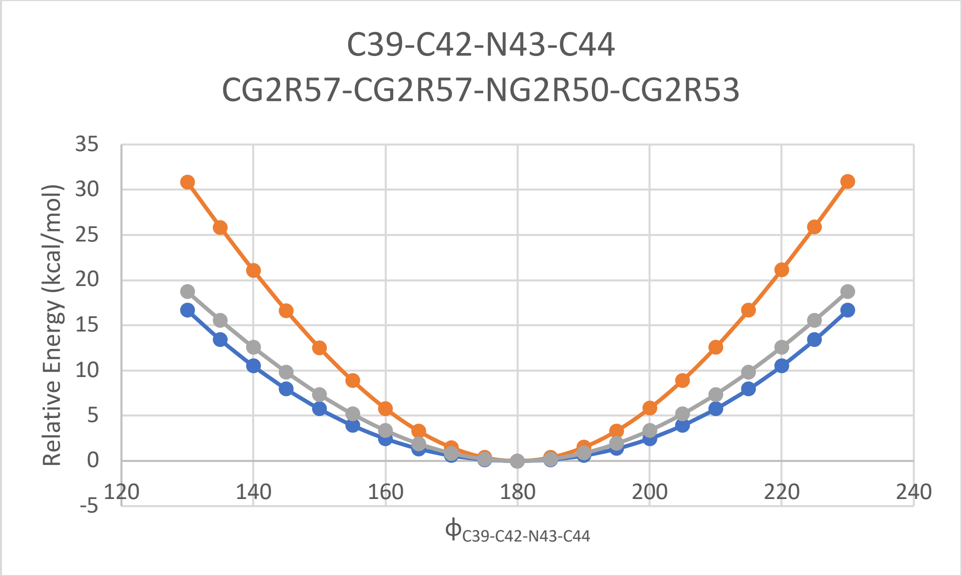


**Supplementary Figure 10.** Potential-energy curve for the C39-C42-N43-C44 dihedral angle in RY785 between 120° and 240°, calculated with MP2/6-31G(d) (blue) and with default CGenFF (orange) or with our optimized forcefield (gray). The Y axis shows the energy (in kcal/mol) relative to the lowest-energy conformer.

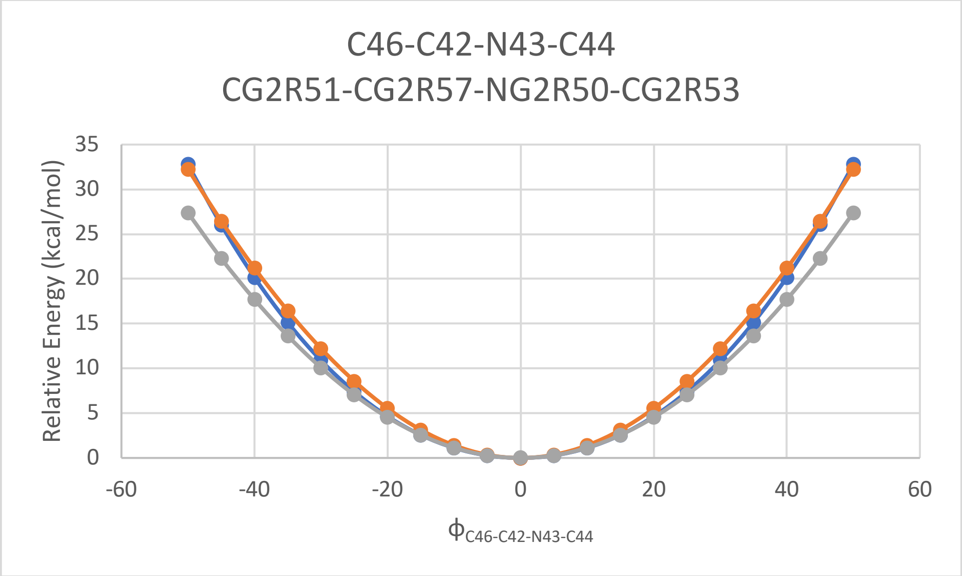


**Supplementary Figure 11.** Potential-energy curve for the C46-C42-N43-C44 dihedral angle in RY785 between -60° and 60°, calculated with MP2/6-31G(d) (blue) and with default CGenFF (orange) or with our optimized forcefield (gray). The Y axis shows the energy (in kcal/mol) relative to the lowest-energy conformer.

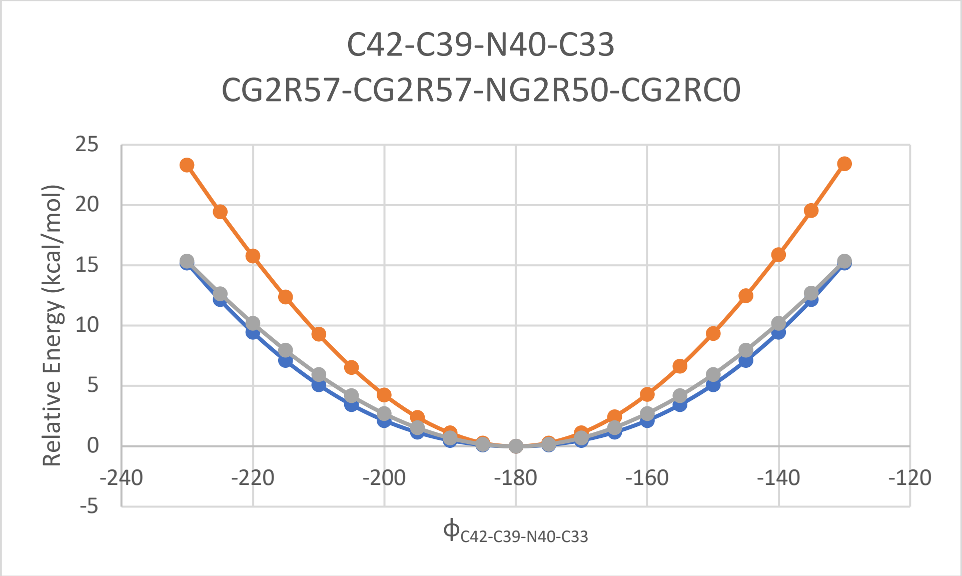


**Supplementary Figure 12.** Potential-energy curve for the C42-C39-N40-C33 dihedral angle in RY785 between -240° and -120°, calculated with MP2/6-31G(d) (blue) and with default CGenFF (orange) or with our optimized forcefield (gray). The Y axis shows the energy (in kcal/mol) relative to the lowest-energy conformer.

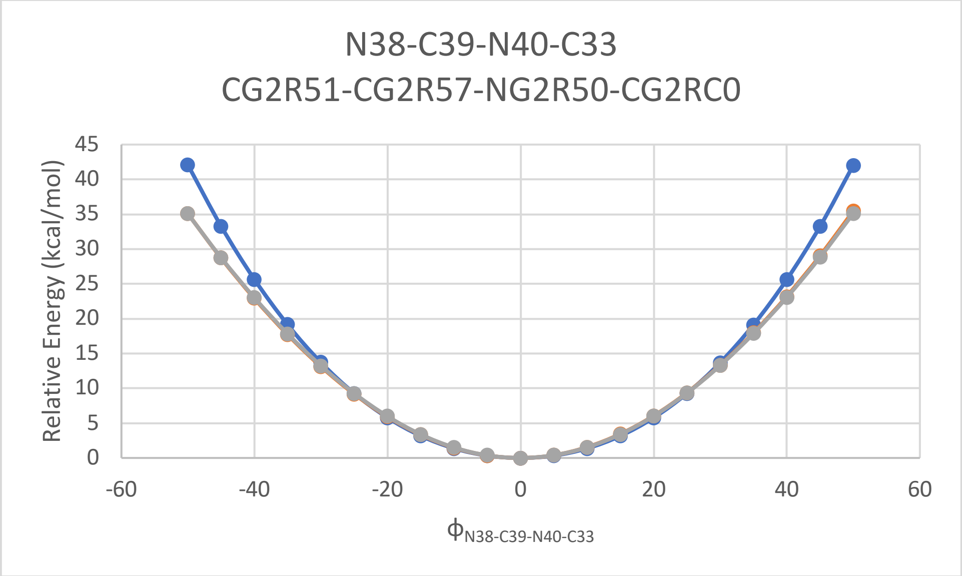


**Supplementary Figure 13.** Potential-energy curve for the N38-C39-N40-C33 dihedral angle in RY785 between -60° and 60°, calculated with MP2/6-31G(d) (blue) and with default CGenFF (orange) or with our optimized forcefield (gray). The Y axis shows the energy (in kcal/mol) relative to the lowest-energy conformer.

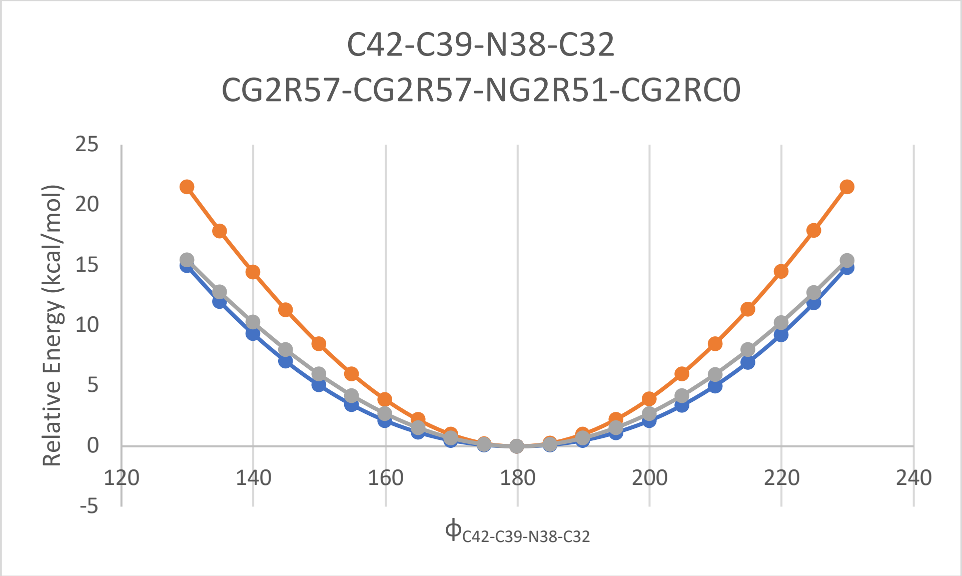


**Supplementary Figure 14.** Potential-energy curve for the C42-C39-N38-C32 dihedral angle in RY785 between 120° and 240°, calculated with MP2/6-31G(d) (blue) and with default CGenFF (orange) or with our optimized forcefield (gray). The Y axis shows the energy (in kcal/mol) relative to the lowest-energy conformer.

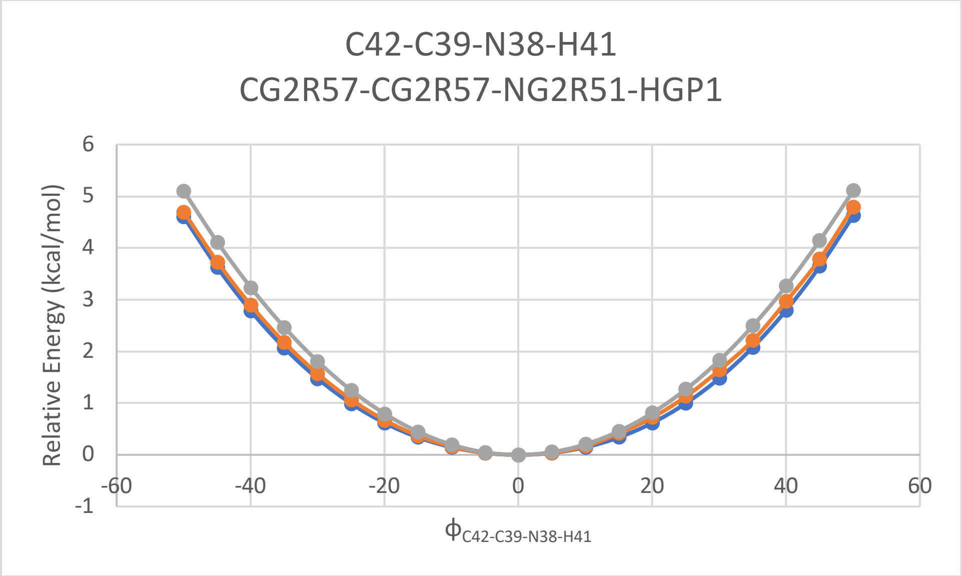


**Supplementary Figure 15.** Potential-energy curve for the C42-C39-N38-H41 dihedral angle in RY785 between -60° and 60°, calculated with MP2/6-31G(d) (blue) and with default CGenFF (orange) or with our optimized forcefield (gray). The Y axis shows the energy (in kcal/mol) relative to the lowest-energy conformer.

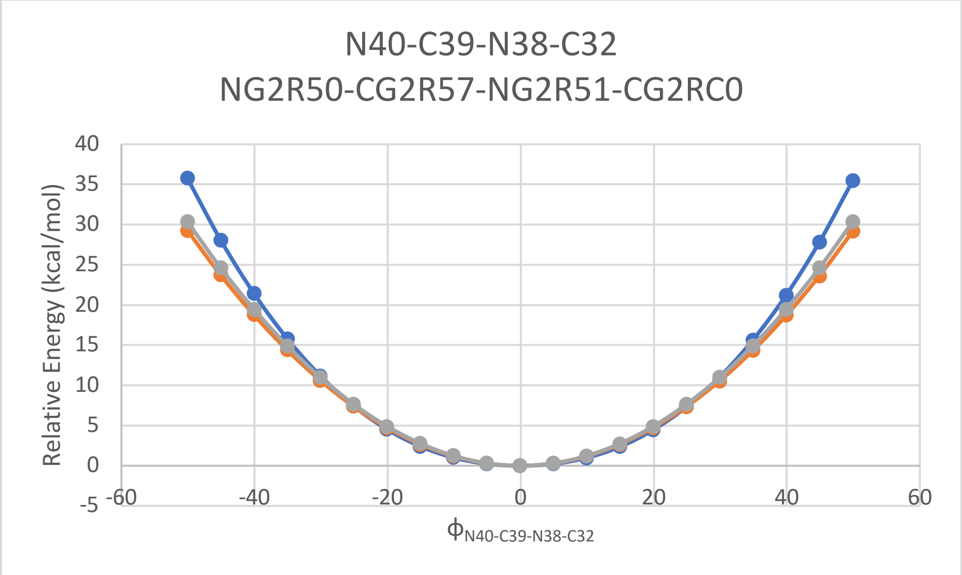


**Supplementary Figure 16.** Potential-energy curve for the N40-C39-N38-C32 dihedral angle in RY785 between -60° and 60°, calculated with MP2/6-31G(d) (blue) and with default CGenFF (orange) or with our optimized forcefield (gray). The Y axis shows the energy (in kcal/mol) relative to the lowest-energy conformer.

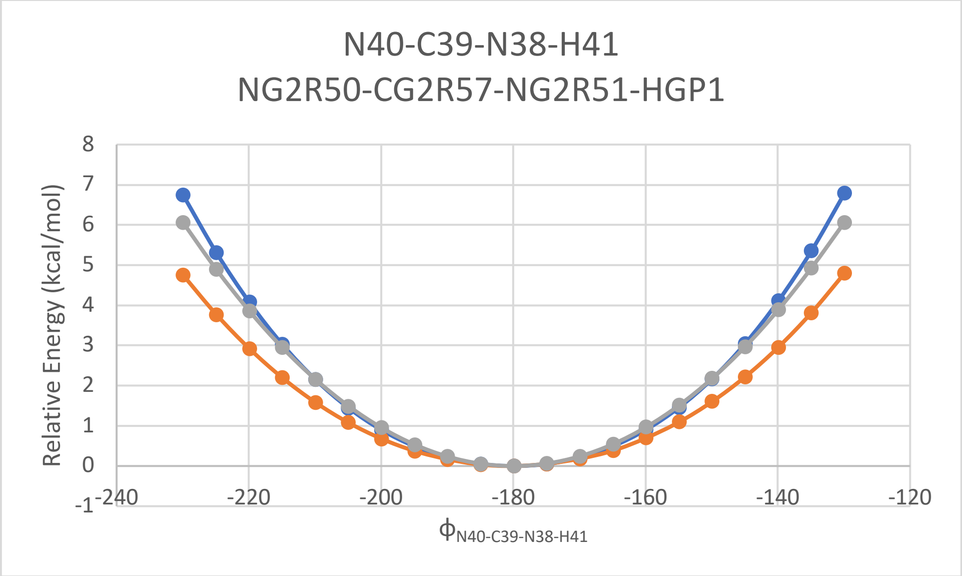


**Supplementary Figure 17.** Potential-energy curve for the N40-C39-N38-H41 dihedral angle in RY785 between -240° and -120°, calculated with MP2/6-31G(d) (blue) and with default CGenFF (orange) or with our optimized forcefield (gray). The Y axis shows the energy (in kcal/mol) relative to the lowest-energy conformer.

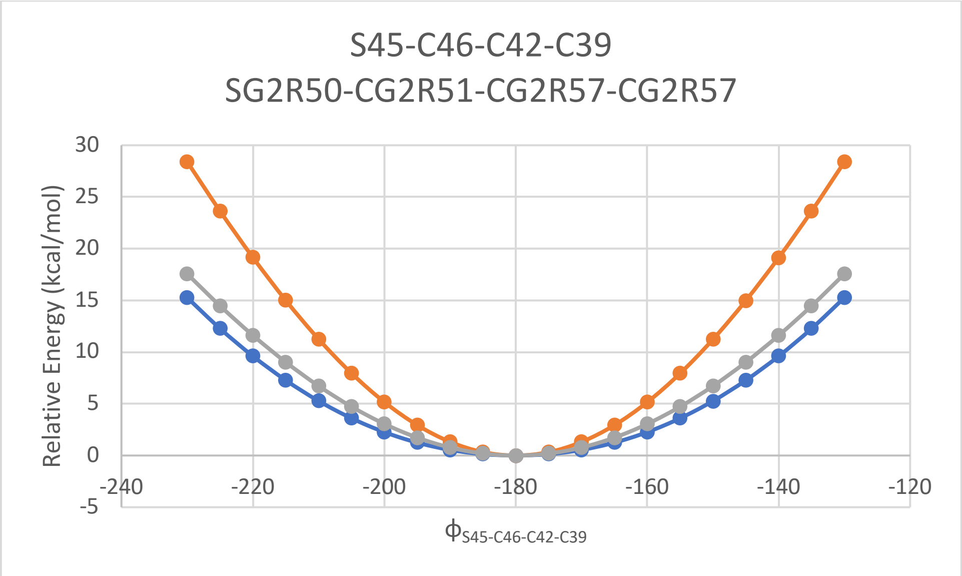


**Supplementary Figure 18.** Potential-energy curve for the S45-C46-C42-C39 dihedral angle in RY785 between -240° and -120°, calculated with MP2/6-31G(d) (blue) and with default CGenFF (orange) or with our optimized forcefield (gray) by scanning. The Y axis shows the energy (in kcal/mol) relative to the lowest-energy conformer.

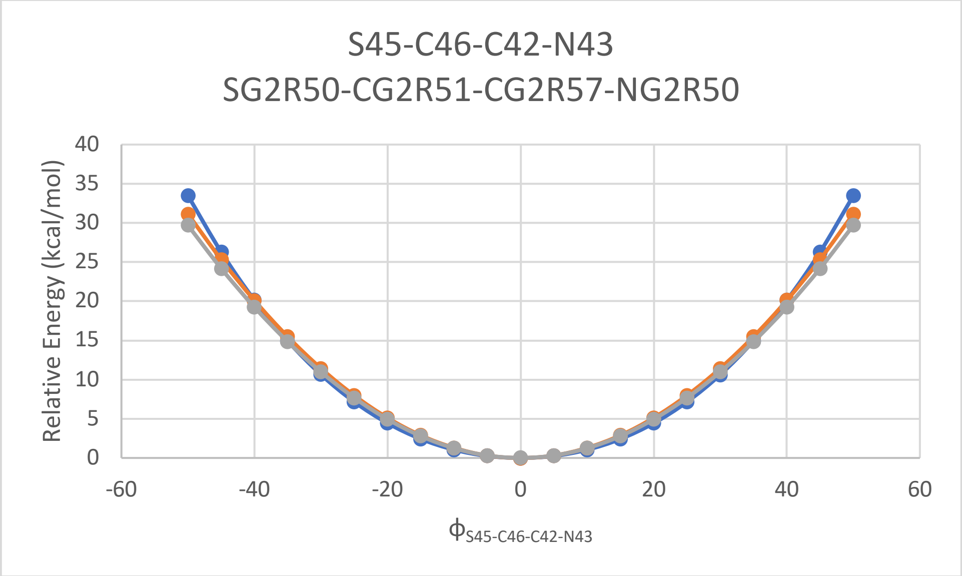


**Supplementary Figure 19.** Potential-energy curves for the S45-C46-C42-N43 dihedral angle in RY785 between -60° and 60°, calculated with MP2/6-31G(d) (blue) and with default CGenFF (orange) or with our optimized forcefield (gray). The Y axis shows the energy (in kcal/mol) relative to the lowest-energy conformer.

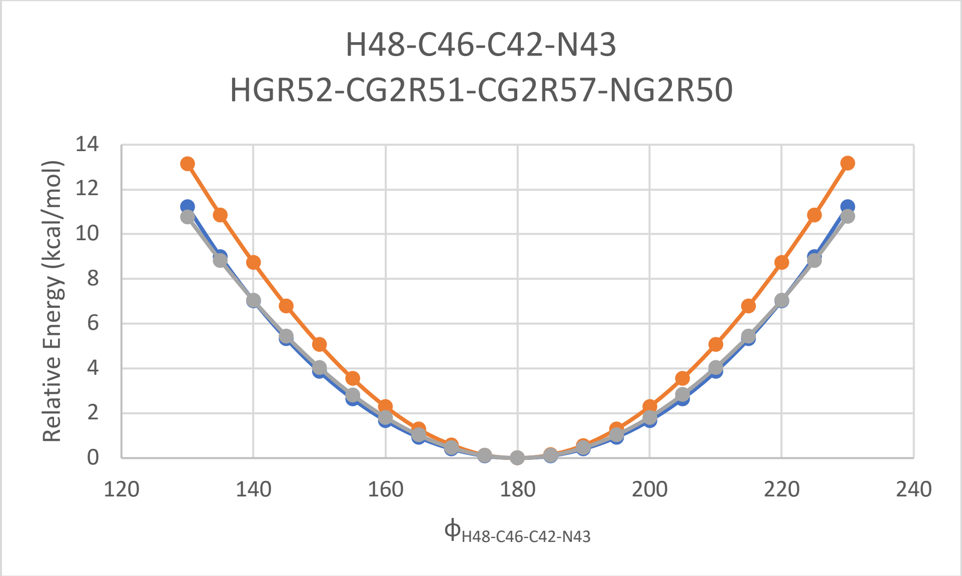


**Supplementary Figure 20.** Potential-energy curve for the H48-C46-C42-N43 dihedral angle in RY785 between 120° and 240°, calculated with MP2/6-31G(d) (blue) and with default CGenFF (orange) or with our optimized forcefield (gray). The Y axis shows the energy (in kcal/mol) relative to the lowest-energy conformer.

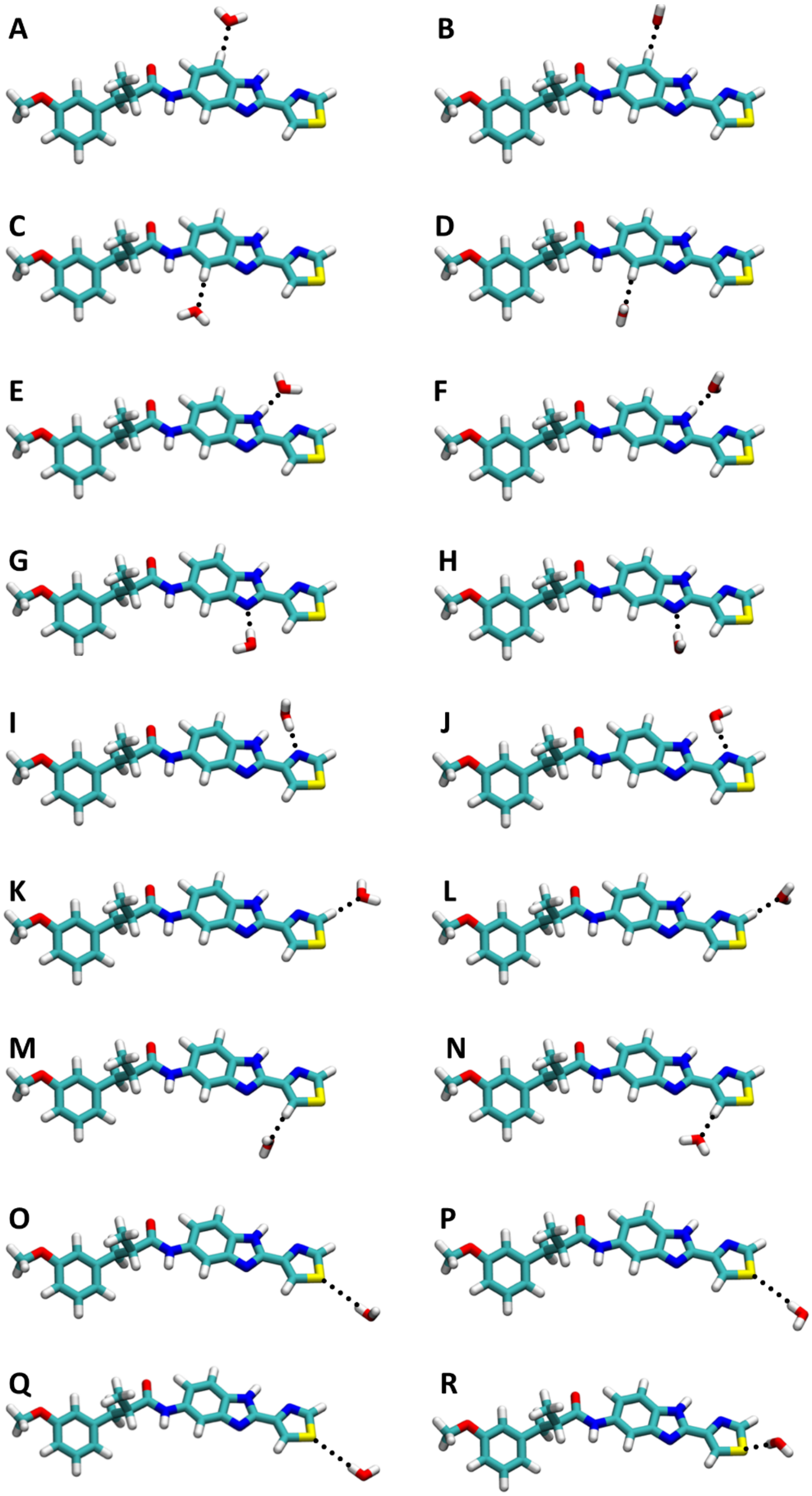


**Supplementary Figure 21.** Ab-initio optimized geometries of RY785-water complexes used in the calibration of our molecular-mechanics forcefield for RY785.

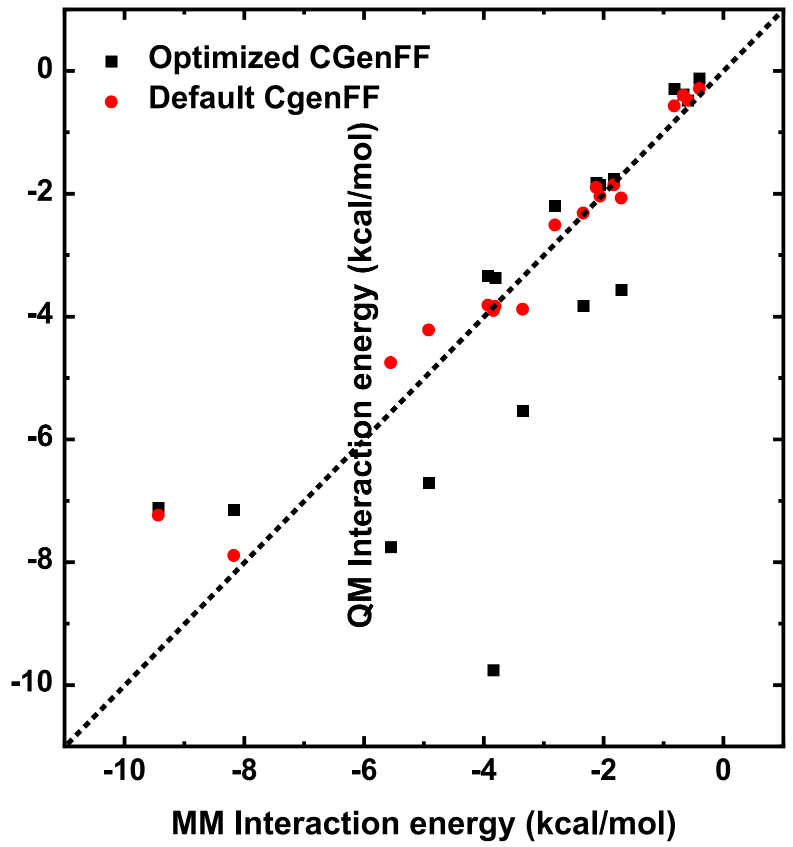


**Supplementary Figure 22.** Interaction energies (in kcal/mol) for the RY785-water complexes shown in Fig. S21. QM data (Y-axis) are correlated with MM values calculated with default CGenFF (red) and with our optimized forcefield (black). The QM interaction energies are calculated with HF/6-31G(d) and scaled by 1.16. The dashed line represents the equation QM interaction energy = MM interaction energy. The default CGenFF for RY785 results in an average unsigned error of 1.2 kcal/mol as compared to an average unsigned error of 0.3 kcal/mol for the optimized model

**Supplementary Table 1.** Interaction distances (in Å) and Interaction energies (in kcal/mol) for the RY785-water complexes shown in Fig. S21. QM data are compared with MM values calculated with default CGenFF and with our optimized forcefield.

| Complex | $E_{QM}$ | $r_{QM}$ | $E_{MM, def}$ | $r_{MM, def}$ | $E_{MM, opt}$ | $r_{MM, opt}$ |
|---|---|---|---|---|---|---|
| **A** | -1.83 | 2.58 | -1.76 | 2.60 | -1.86 | 2.59 |
| **B** | -2.12 | 2.51 | -1.82 | 2.60 | -1.90 | 2.60 |
| **C** | -1.71 | 2.61 | -3.56 | 2.53 | -2.07 | 2.64 |
| **D** | -2.34 | 2.47 | -3.83 | 2.51 | -2.32 | 2.62 |
| **E** | -4.92 | 1.99 | -6.70 | 1.83 | -4.22 | 1.92 |
| **F** | -3.35 | 2.14 | -5.53 | 1.87 | -3.88 | 1.94 |
| **G** | -8.17 | 2.06 | -7.15 | 1.88 | -7.89 | 1.87 |
| **H** | -9.44 | 2.04 | -7.11 | 1.88 | -7.24 | 1.88 |
| **I** | -5.56 | 2.09 | -7.75 | 1.85 | -3.90 | 1.97 |
| **J** | -3.84 | 2.07 | -9.76 | 1.83 | -4.75 | 1.95 |
| **K** | -3.81 | 2.32 | -3.38 | 2.23 | -3.84 | 2.22 |
| **L** | -3.93 | 2.30 | -3.34 | 2.24 | -3.82 | 2.22 |
| **M** | -2.06 | 2.31 | -1.86 | 2.26 | -2.04 | 2.25 |
| **N** | -2.81 | 2.22 | -2.20 | 2.22 | -2.51 | 2.21 |
| **O** | -0.67 | 2.86 | -0.38 | 2.60 | -0.40 | 2.62 |
| **P** | -0.59 | 2.92 | -0.49 | 2.59 | -0.48 | 2.60 |
| **Q** | -0.40 | 2.94 | -0.12 | 2.66 | -0.29 | 2.67 |
| **R** | -0.82 | 2.80 | -0.29 | 2.58 | -0.37 | 2.57 |
